# T-Cell-Mediated Killing in Glioblastoma Organoids induced by Adenoviral Delivery of the CIITA Transgene

**DOI:** 10.1101/2024.01.22.576747

**Authors:** Ilaria Salvato, Eliane Klein, Aurélie Poli, Mahsa Rezaeipour, Luca Ermini, Bakhtiyor Nosirov, Anuja Lipsa, Anais Oudin, Virginie Baus, Gian Mario Dore, Antonio Cosma, Anna Golebiewska, Antonio Marchini, Simone P. Niclou

## Abstract

The immunosuppressive nature of the tumor microenvironment poses a significant challenge to effective immunotherapies against glioblastoma (GB). Boosting the immune response is critical for a successful therapy. Here, we adopted a cancer gene therapy approach to induce T-cell mediated killing of the tumor through increased activation of the immune system. Patient-based 3D GB models were infected with a replication-deficient adenovirus (AdV) armed with the Class II Major Histocompatibility Complex (MHC-II) Transactivator *CIITA* gene (Ad-CIITA). Successful induction of surface MHC-II was achieved in infected GB cell lines and primary human GB organoids. Infection with an AdV carrying a mutant form of *CIITA* with a single amino acid substitution resulted in cytoplasmic accumulation of CIITA without subsequent MHC-II expression. Co-culture of infected tumor cells with either PBMCs or isolated T-cells led to dramatic breakdown of GB organoids. Intriguingly, both wild-type and mutant Ad-CIITA but not unarmed AdV, triggered immune-mediated tumor cell death in the co-culture system, suggesting an at least partially MHC-II-independent process. We further show that the observed cancer cell killing requires the presence of either CD8^+^ or CD4^+^ T-cells and the direct contact between GB and immune cells. We did not however detect evidence of activation of canonical T-cell mediated cell death pathways. While the precise mechanism remains to be determined, these findings highlight the potential of AdV-mediated *CIITA* delivery to enhance T-cell-mediated immunity against GB.

## INTRODUCTION

Malignant brain tumors including glioblastoma (GB) remain a major clinical and scientific challenge. The immunosuppressive tumor microenvironment (TME) as well as systemic immunosuppression pose challenges to current immunotherapies. GBs recruit immunosuppressive cells like tumor-associated microglia and macrophages (TAMs) and regulatory T-cells (Tregs), while only a limited number of T-cells are present, often showing signs of exhaustion^1–3^. GB tumors may also downregulate the expression of major histocompatibility complex (MHC) molecules that play a central role in presenting cancer-derived antigens to the immune system^4^.

Despite promising results in preclinical studies, immunotherapies, whether tested individually or in combination with standard care, have so far failed to translate in clinically meaningful outcomes for GB patients ^5^. This discrepancy can be largely attributed to oversimplified preclinical models that do not recapitulate the complex biology of GBs. Replication-deficient adenoviruses (AdVs) have demonstrated safety and tolerability in the treatment of various cancer types, including GB. Current GB treatments using AdVs focus on the delivery of herpes simplex virus (HSV) thymidine kinase (TK), p53, interleukin (IL)-12, interferon (IFN)-β, and a chimeric death receptor (VB-111)^5^.

A promising approach is to arm non-lytic AdV vectors with transgenes to boost anti-GB immunity. Class II Major Histocompatibility Complex Transactivator (CIITA) is the master regulator of MHC class II (MHC-II) molecules, typically expressed in professional antigen presenting cells (APCs)^6^. Upon expression and translocation to the nucleus, CIITA interacts with the constitutively-expressed factors of the enhanceosome complex and activates the transcription of cell surface “classical” human leukocyte antigen (HLA)-DR, HLA-DP and HLA-DQ, which present antigens to CD4^+^ T-cells, as well as HLA-DM, HLA-DO and invariant chain (Ii), involved in intracellular antigen processing^7^.

Previous work on carcinoma cell lines^8–10^, and more recently on murine glioma cells^11^, found that ectopic stable expression of CIITA transforms tumor cells into MHC-II-positive cells with enhanced antigen processing and presentation capabilities. Remarkably, CD80 and CD86 costimulatory molecules, essential alongside MHC-II expression, were discovered to be dispensable for T-cell activation^8, 12, 13^. In these models, CIITA-transfected/infected MHC-II-positive cells were found to prime and activate CD4^+^ T-cells, resulting in tumor rejection or growth retardation in mice with subcutaneous tumors^10–13^. However, direct antigen presentation and antigen-based T-cell activation have not been reported. A recent study indicated limited pro-immunogenic potential of CIITA in GB tumors^14^.

Here, we engineered a replication-deficient AdV with *CIITA* to infect patient-derived GB organoids and promote antitumor immunity. We established an advanced human GB co-culture system, based on primary organoids co-cultured with human immune cells, to assess the functional impact of the gene therapy approach in a complex model. We find that AdV-mediated delivery of *CIITA* induces cancer cell death in the presence of immune cells. While the exact mechanism remains to be elucidated, our data suggest that the effect is at least partially independent of MHC-II expression.

## RESULTS

### MHC-II expression in GB is limited to non-neoplastic cells of the tumor microenvironment

To assess the diversity of MHC-II molecules in GB, we examined gene expression profiles at bulk and single-cell level in patient tumors and preclinical models. At bulk RNA level, expression of *CIITA* and genes coding for MHC-II antigen presenting molecules (*HLA-DQA1*, *HLA-DQB1*, *HLA-DRA, HLA-DRB1, HLA-DRB5)* was higher in GBs than in normal brain tissue (**Fig.1A**). However, analysis of single-cell RNA-sequencing (RNA-seq) of GB patient tumors revealed that tumor cells do not express *CIITA* nor genes encoding for MHC-II antigen processing (*CD74*, *HLA-DMA*, *HLA-DOA*) and presentation (*HLA-DQA1*, *HLA-DQB1*, *HLA-DRA, HLA-DRB1, HLA-DRB5)* (**Fig.1B**). High expression of these genes was detected in immune cells of the TME, explaining the high transcript levels observed by bulk RNA-seq. The highest MHC-II expression was detected in well-known APCs, including dendritic cells, macrophages, microglia, and B cells (**Fig.1B**). Limited cancer cell expression of MHC-II genes was confirmed in a cohort of 28 GB patient-derived orthotopic xenografts (PDOXs), where bulk RNA-seq profiles showed low expression of human genes involved in MHC-II antigen processing and presentation and moderately high levels of enhanceosome-associated genes (**Fig.1C**). Except for U87 cell line, similar expression profiles were seen in human GB cell lines, including adherent and 3D sphere cultures of GB stem-like cells (GSCs) (**Supplementary Fig.1A**). Interestingly, the genes associated with the MHC-II enhanceosome complex (*CREB1*, *NFYA*, *NFYB*, *NFYC*, *RFX5*, *RFXANK*, and *RFXAP)* showed sustained expression throughout patient tumors, PDOX and cell lines (**Supplementary Fig.1A**), suggesting that GB cells have the capacity to express MHC-II upon activation of the MHC-II master regulator CIITA. Moreover, most of the preclinical models exhibited transcriptional expression of the Coxsackie and Adenovirus Receptor (*CXADR*), the specific receptor for AdV serotype 5, as well as the integrin αvβ5 necessary for AdV infection (*ITGAV* and *ITGB5*)^15^ (**Fig.1C, Supplementary Fig.1A**). This observation implies that GB tumor cells are likely to be susceptible to AdV infection, highlighting the suitability of non-lytic AdVs for targeting GB.

**Figure 1.**
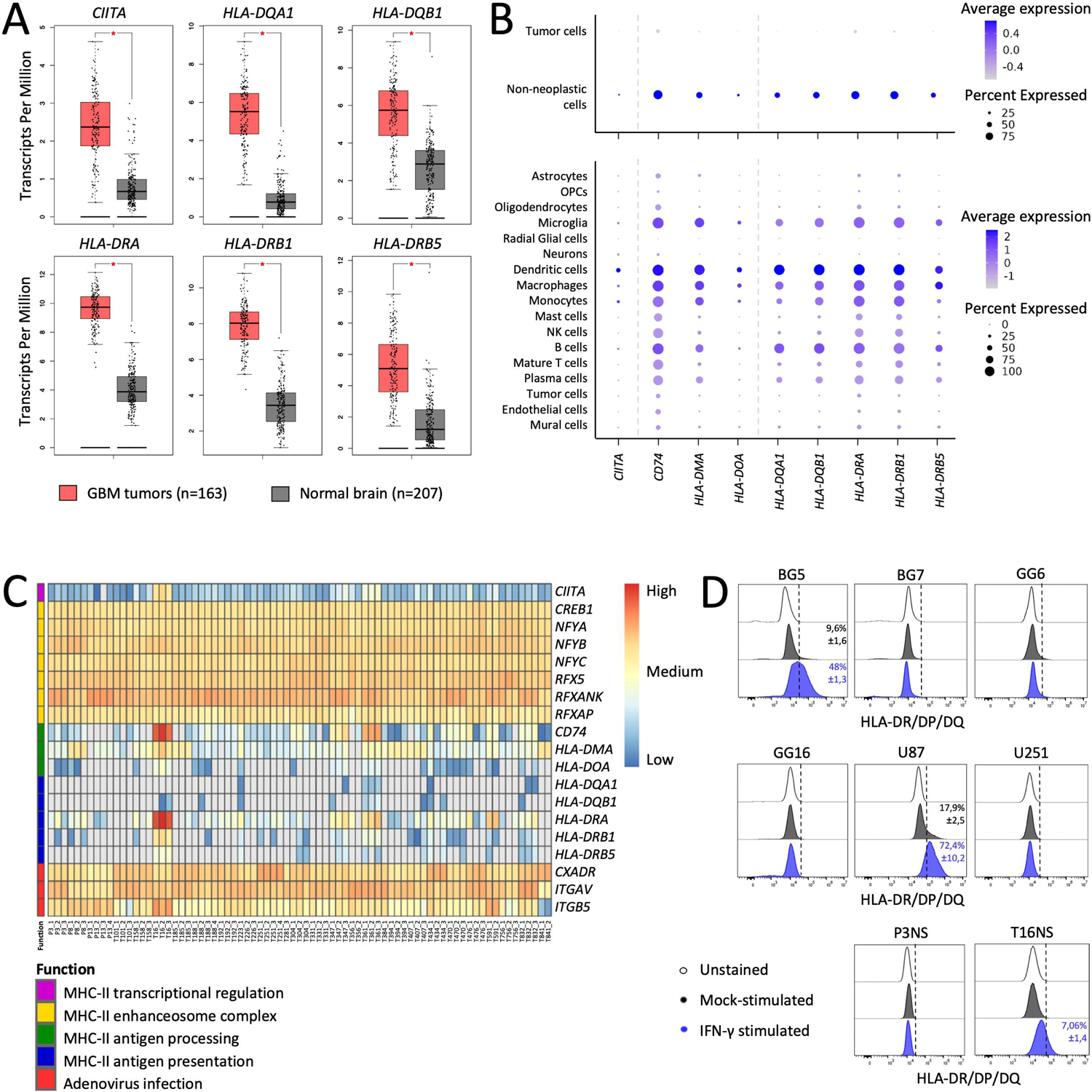
Expression of *MHC-II*-related transcripts in human glioblastoma and associated models. **A**. Bulk RNA-seq gene expression in GB patient tumors (n=163) versus control normal brain (n=207). *p<0.01 (one-way ANOVA). **B.** Single-cell RNA-seq gene expression in GB patient tumors. **C.** Heatmap representing bulk RNA-seq (human specific) gene expression in GB PDOXs. Grey color indicates lack of expression. PDOX replicates are indicated by numbers. **D.** FACS analysis of surface MHC-II (anti-HLA-DR/DP/DQ-FITC), in control conditions (cells unstained or mock-stimulated) and upon IFN-γ stimulation.

Next, we assessed the presence of MHC-II proteins at the cell surface of GB cells by flow cytometry. The majority of GB cells showed no MHC-II protein at the cell surface under regular culture conditions (**Fig.1D, Supplementary Fig.1B**). Out of 10 GB cell lines analyzed, only U87 and BG5 cells showed some positive cells (17.9% and 9.6% respectively). Interestingly, IFN-γ cytokine stimulus did not activate MHC-II expression in GB cells, except for U87, BG5, and T16NS (**Fig.1D**).

In summary, we confirmed that MHC-II molecules are largely absent from GB tumor cells but are expressed in tumor-associated APCs. Based on the presence of transcripts of the MHC-II enhanceosome complex, ectopic expression of CIITA appears as a valid strategy to deliver MHC-II molecules at the cell membrane. Moreover, the expression of the necessary AdV receptor and cell adhesion molecules confers infectivity to GB cells to serotype 5 AdVs.

### Human GB stem-like cells infected with Ad-CIITA express CIITA and present MHC-II molecules at the cell surface

To convert MHC-II-negative GB tumor cells into APC-like cells, we generated a replication-deficient AdV carrying the wild-type *CIITA* gene (Ad-CIITA) (**Fig.2A**) or a mutated *CIITA* transgene (Ad-CIITA mutant). The mutant CIITA carried a novel missense point mutation (guanine-to-adenosine) (**Supplementary Fig.2A**) leading to a single amino acid substitution (glycine (G) to arginine (A)) at position 1088 (**Supplementary Fig.2B**). Serendipitously generated by us, we found that this mutation impaired the translocation of CIITA to the cell nucleus (**Supplementary Fig.2C**). AdV vectors without transgene (Ad-null) or with the green fluorescent protein (GFP)-encoding transgene (Ad-GFP) were used as controls (**Fig.2A**).

**Figure 2.**
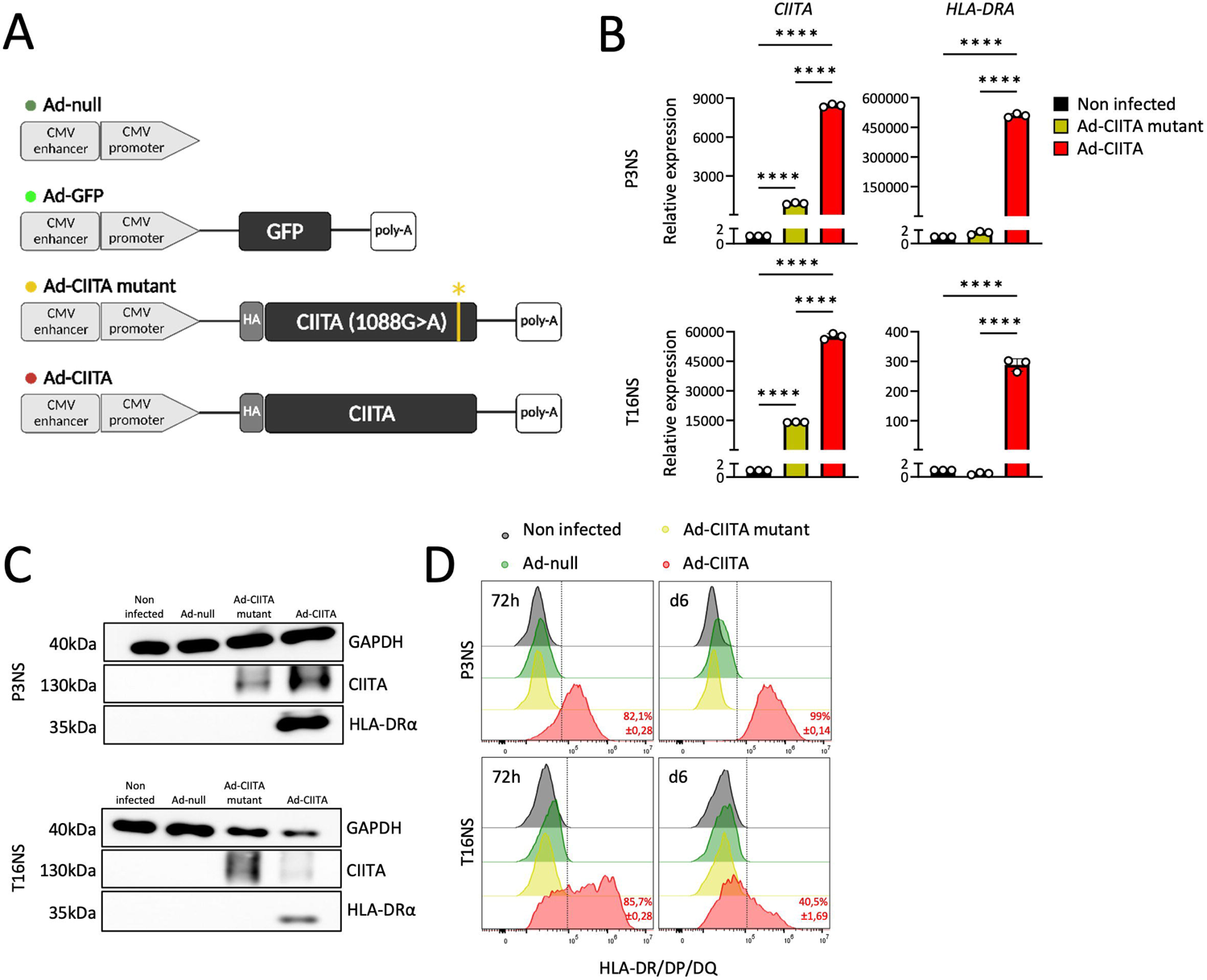
Characterization of Ad-CIITA and Ad-CIITA mutant in human glioblastoma stem-like cells. **A**. AdV vectors used in the study. **B-D**. Virus quality check in human P3NS and T16NS infected at MOI 50. mRNA (**B**) and protein levels (**C**) of CIITA and HLA-DRα were analyzed 72 hours post-infection by qRT‒PCR and Western blot, respectively. qRT‒PCR was normalized to Ef1a, while Western blot to GAPDH. Mean ± SD. ****p<0.0001 (one-way ANOVA). **D**. FACS analysis of surface MHC-II (anti-HLA-DR/DP/DQ-FITC) at 72 hours and 6 days post-infection.

Optimal viral infection and appropriate CIITA and MHC-II expression were determined in the mouse GB cell line GL261 (**Supplementary Fig.2D-E**). In human GSC lines (P3NS and T16NS), CIITA mRNA (**Fig.2B**) and protein (**Fig.2C**) were detected at 72 hours post-infection with both Ad-CIITA and Ad-CIITA mutant. Only wild-type CIITA induced HLA-DRα mRNA and protein expression (**Fig.2B-C**), the latter was confirmed at the cell membrane (**Fig.2D**). In summary, we successfully generated an Ad-CIITA vector that led to surface MHC-II expression in human GB cells. Ad-CIITA mutant vector did not induce MHC-II expression allowing to distinguish between nuclear and non-nuclear CIITA effects.

### Infection with Ad-CIITA induces MHC-II expression in human primary GB organoids

We next determined the efficacy of Ad-CIITA in primary GB organoids from different patients (P3, T16 and T188), representing tumors with classical genetic and epigenetic features (**Table S1**). In P3 organoids, a viral multiplicity of infection (MOI) between 25 and 100 led to strong ectopic expression of CIITA and HLA-DRα at 72-post-infection (**Fig.3A**), whithout a major effect on tissue structure (**Fig.3B**). Again, only Ad-CIITA-infected cells showed MHC-II expression (**Fig.3B**, lower panel), also confirmed by flow cytometry (**Fig.3C**). MHC-I (HLA-A/B/C) expression remained largely unaltered, while CD80 or CD86 co-stimulatory molecules were undetectable at baseline or upon infection (**Supplementary Fig.3A**).

**Figure 3.**
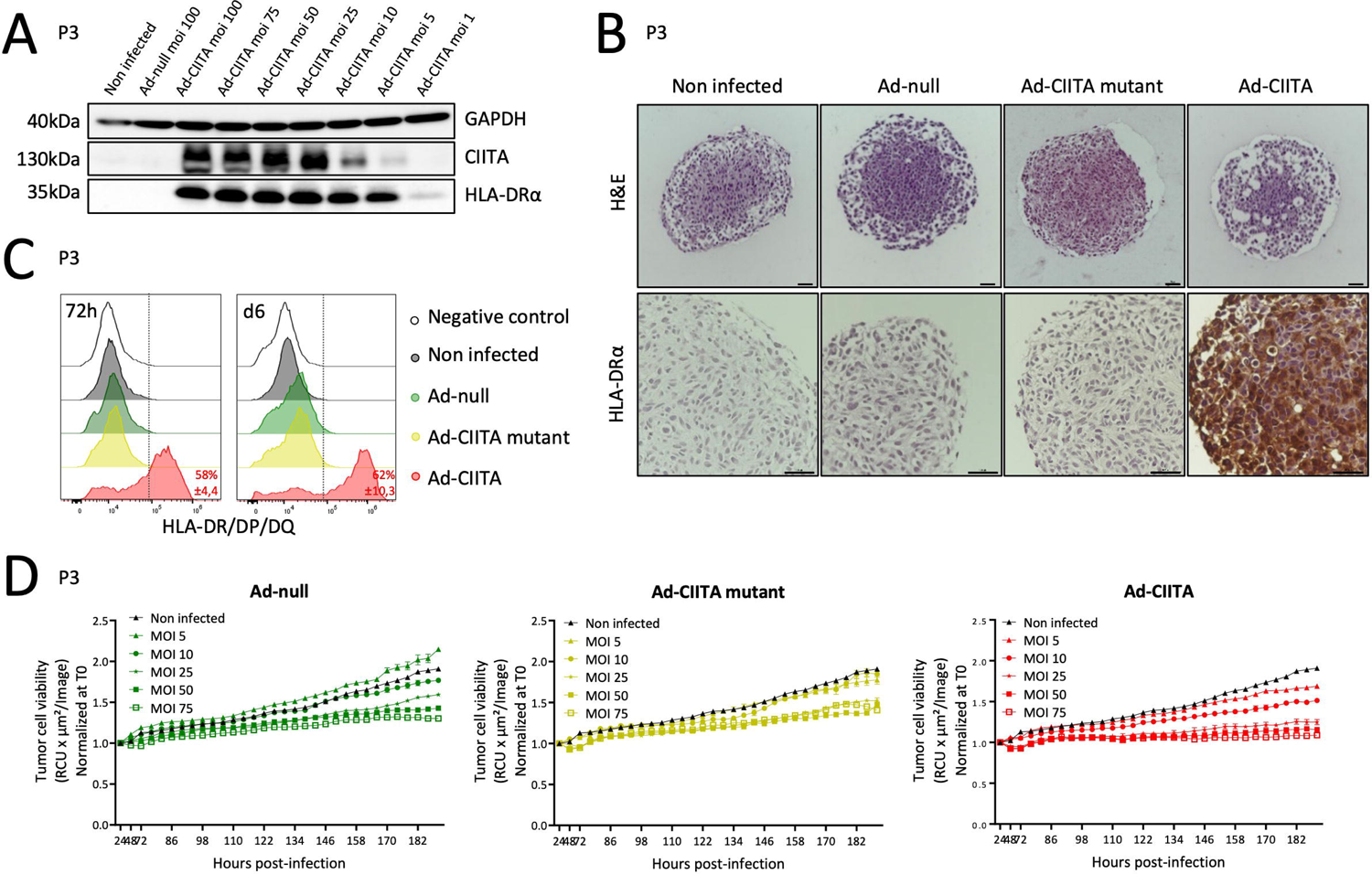
Successful infection of human primary glioblastoma organoids by adenoviral vectors. **A.** Western Blot analysis of CIITA and HLA-DRα expression at 72 hours post-infection in P3 organoids. **B.** Representative H&E (top) and anti-HLA-DRα (bottom) staining at 72 hours post-infection (MOI 50). **C.** FACS analysis of surface MHC-II (anti-HLA-DR/DP/DQ-FITC) at 72 hours and 6-days post-infection (MOI 50). **D.** Evaluation of virus-mediated cytotoxicity (P3dsRed) via Incucyte S3. Mean ± SEM.

To evaluate potential adverse effects of AdV, dsRed-expressing P3 organoids were infected at increasing MOIs, and dsRed fluorescence intensity was monitored in real-time. As expected^16^, viral infection caused a small dose-dependent reduction in organoid growth rate, independent of the transgene, but did not show signs of cell death (**Fig.3D**). H&E-stained sections confirmed this observation (**Fig.3B**). Similar results were seen in T16 organoids, while T188 organoids were unaffected by viral infection (**Supplementary Fig.3B**). In conclusion, AdVs effectively infect primary GB organoids with minimal impact on tumor growth. Robust MHC-II expression is obtained with Ad-CIITA, but not with Ad-CIITA mutant or Ad-null vectors.

### Ad-CIITA-triggered immune-mediated tumor cell death is mediated by CD3^+^ T-cells but is largely independent of MHC-II expression

To evaluate the potential of MHC-II-expressing human GB to elicit adaptive immune responses, we established a protocol involving the co-culture of virally-infected primary GB organoids with allogenic immune cells, from healthy donors or from partially HLA-matched donors (**Fig.4A**). The purity of PBMCs and isolated cell types were assessed by flow cytometry before co-culture (**Supplementary Fig.4A, Table S2**).

**Figure 4.**
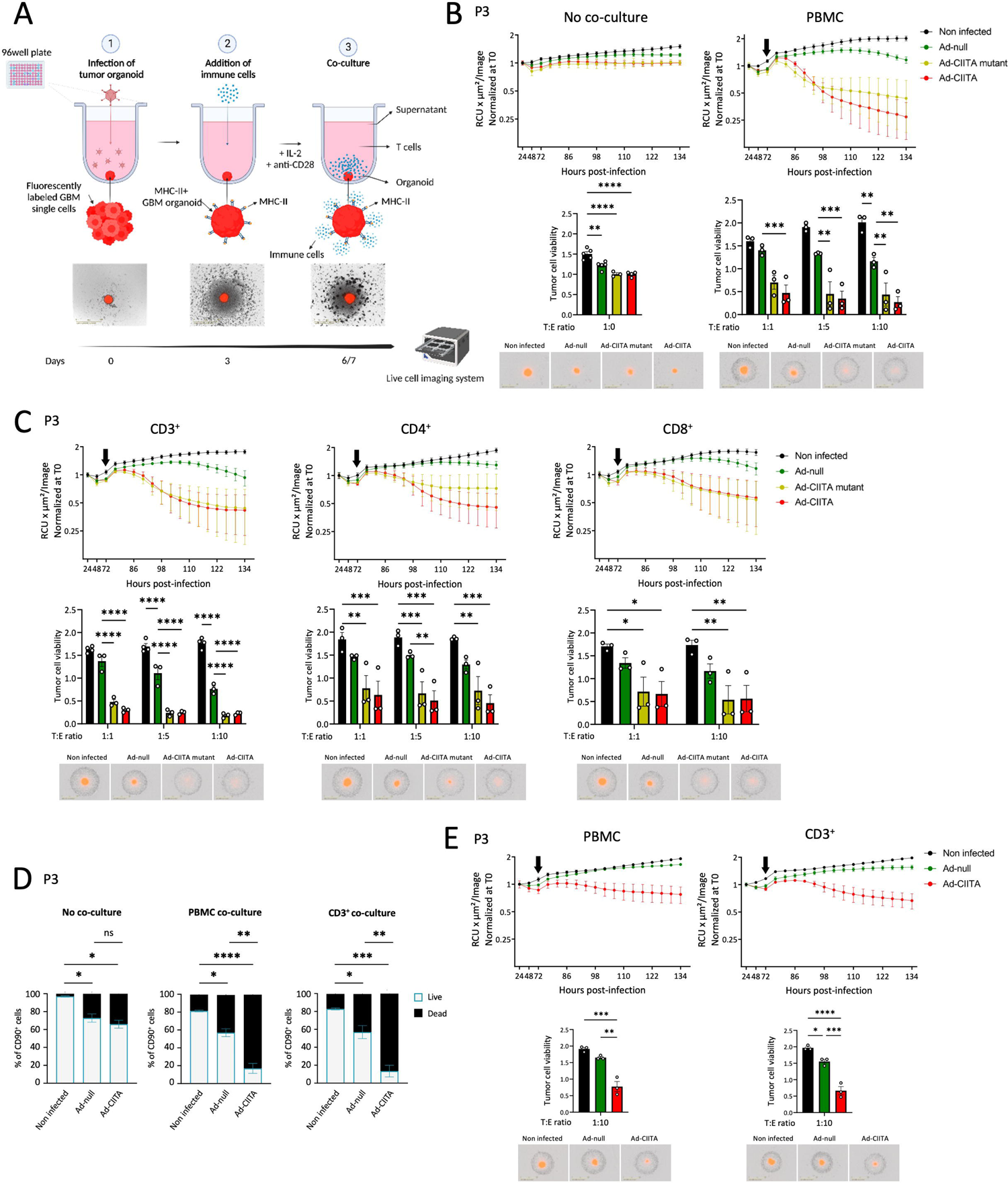
Immune cell-mediated tumor cell killing in human primary GB organoids induced by Ad-CIITA and Ad-CIITA mutant. **A.** Schematic of AdV-infected GB organoids co-cultured with immune cells. **B-C.** Tumor cell viability in P3dsRed organoids (MOI 75): **(B)** alone (no co-culture) or co-cultured with PBMCs; **(C)** co-cultured with CD3^+^, CD4^+^ or CD8^+^ T-cells; Mean ± SEM. *p<0.05, **p<0.01, ***p<0.001, and ****p<0.0001 (two-way ANOVA). **D.** FACS analysis showing the proportion of live/dead cells in CD90^+^ tumor cells at 72 hours post-co-culture: alone (left), co-cultured with PBMCs (middle), and co-cultured with CD3+ T-cells (right). Mean ± SD. *p<0.05, **p<0.01, ***p<0.001, and ****p<0.0001 (one-way ANOVA). **E.** Tumor cell viability in P3dsRed organoids (MOI 75) co-cultured with HLA-matched PBMCs or CD3^+^ T-cells. Mean ± SEM. *p<0.05, **p<0.01, ***p<0.001, and ****p<0.0001 (two-way ANOVA).

Based on previous tumor-immune cell co-culture studies^17^, we first used immune cells from healthy human donors. In co-culture conditions without immune cells, P3 GB organoids retained their structure, their proliferation being slightly affected by viral infection (**Fig.4B**), in line with the experiments shown in **Fig.3D**. Upon addition of allogenic PBMCs, we observed a pronounced organoid disruption and tumor cell killing in Ad-CIITA-infected organoids (86,5%±10,2), contrasting with a lower effect in Ad-null-infected organoids (42,2%±7,3) (**Fig.4B, Video S1**). Percentages represent the reduction in dsRed signal compared to non-infected controls at the co-culture end-point. Interestingly, the Ad-CIITA mutant also led to robust tumor cell killing in the presence of PBMCs (78,3%±21,7) (**Fig.4B, Video S1**), despite the lack of MHC-II expression (**Fig.3C**). Similar results were seen in T16 and T188 organoids, where pronounced tumor cell death was induced by Ad-CIITA and again, though to a lesser extent, by Ad-CIITA mutant (**Supplementary Fig.4B**). These data suggest that CIITA expression potentiates the immunogenicity of GB organoids for alloreactive PMBCs, potentially mediated by a cumulative effect of MHC-II-dependent and independent mechanisms.

We next sought to determine whether CD3+ T-cells were involved in the observed killing phenotype. Similar to PBMCs, purified CD3^+^ T-cells alone efficiently induced tumor cell killing in P3 organoids infected with Ad-CIITA (76,4%±21,9) or Ad-CIITA mutant (75,1%±29,4), suggesting a primary T-cells involvement (**Fig.4C vs Fig.4B**). Immune-mediated tumor cell death was further confirmed by flow cytometry (**Fig.4D**). To assess the specificity of Ad-CIITA induced T-cell killing, we performed co-culture experiments with purified CD4^+^ and CD8^+^ T-cells, which rely on MHC-II and MHC-I molecules, respectively. Efficient tumor cell killing was maintained with either CD4^+^ (Ad-CIITA: 75,6%±16,8) or CD8^+^ T-cells (Ad-CIITA: 67,5%±28,8) (**Fig.4C**). Similar results were obtained with T16 and T188 organoids (**Supplementary Fig.4C**), reinforcing the involvement of a MHC-II-independent mechanism in GB cell killing. Nevertheless, we next aimed to exclude a purely alloreactive T-cell reaction, by applying effector cells sharing a partial match at HLA alleles with the target GB organoids. Unfortunately, autologous PBMCs from the tumor patient were not available. Specifically, PBMCs were chosen to match the HLA typing of the donor GB tumors at least at one allele of each MHC-I (*HLA-A*, *HLA-B*, *HLA-C*) and each MHC-II (*HLA-DRB1/3/4/5*, *HLA-DQA1*, *HLA-cytokines upon tumor cell killing mediated DQB1*, *HLA-DPA1*, *HLA-DPB1*) locus. The matching degree varied depending on the donor, as presented in **Table S3**. The Ad-CIITA-mediated killing phenotype was retained in the P3 co-culture model (**Fig.4E**) and confirmed in T16 tumors (**Supplementary Fig.4D**). Although the cytotoxic activity of HLA-matched PBMCs and isolated CD3^+^ cells was reduced compared to allogenic PBMCs (**Fig.4C vs Fig.4E**), these data confirmed that the main killing activity observed could not be attributed to an alloreactive immune reaction linked to mismatching MHC molecules.

In summary, our findings show that both Ad-CIITA and Ad-CIITA mutant enhance the immunogenicity of GB for allogenic T-cells, facilitating efficient tumor cell killing in the presence of either CD4+ or CD8+ T-cells. These data suggest that CIITA activates a T-cell-mediated immune response that is at least partially independent of MHC-II antigen presentation.

### Immune-mediated tumor cell killing of GB organoids requires CIITA expression and cell-to-cell contact with T-cells, but no antigen presentation

To address the functionality of MHC-II molecules induced by Ad-CIITA, we used a proliferation assay of T-cells from OT-II mice, specifically reacting to ovalbumin peptides 323-339 (OVA323-339) (**Fig.5A**). In contrast to the positive control, CD3^+^ T-cells co-cultured with Ad-CIITA-infected OVA323-339-pulsed GL261 cells, did not proliferate, indicating that the MHC-II molecules present on infected GL261 cells, did not present antigens to T-cells, reinforcing the MHC-II independence of the observed phenotype. This was further confirmed in the human GB organoid - T-cell co-cultures in the presence of an antigenic pool consisting of cytomegalo-, parainfluenza, and influenza virus peptides (“CPI pool”). No difference in tumor cell death was observed between pulsed and unpulsed co-cultures (**Fig.5B**).

**Figure 5.**
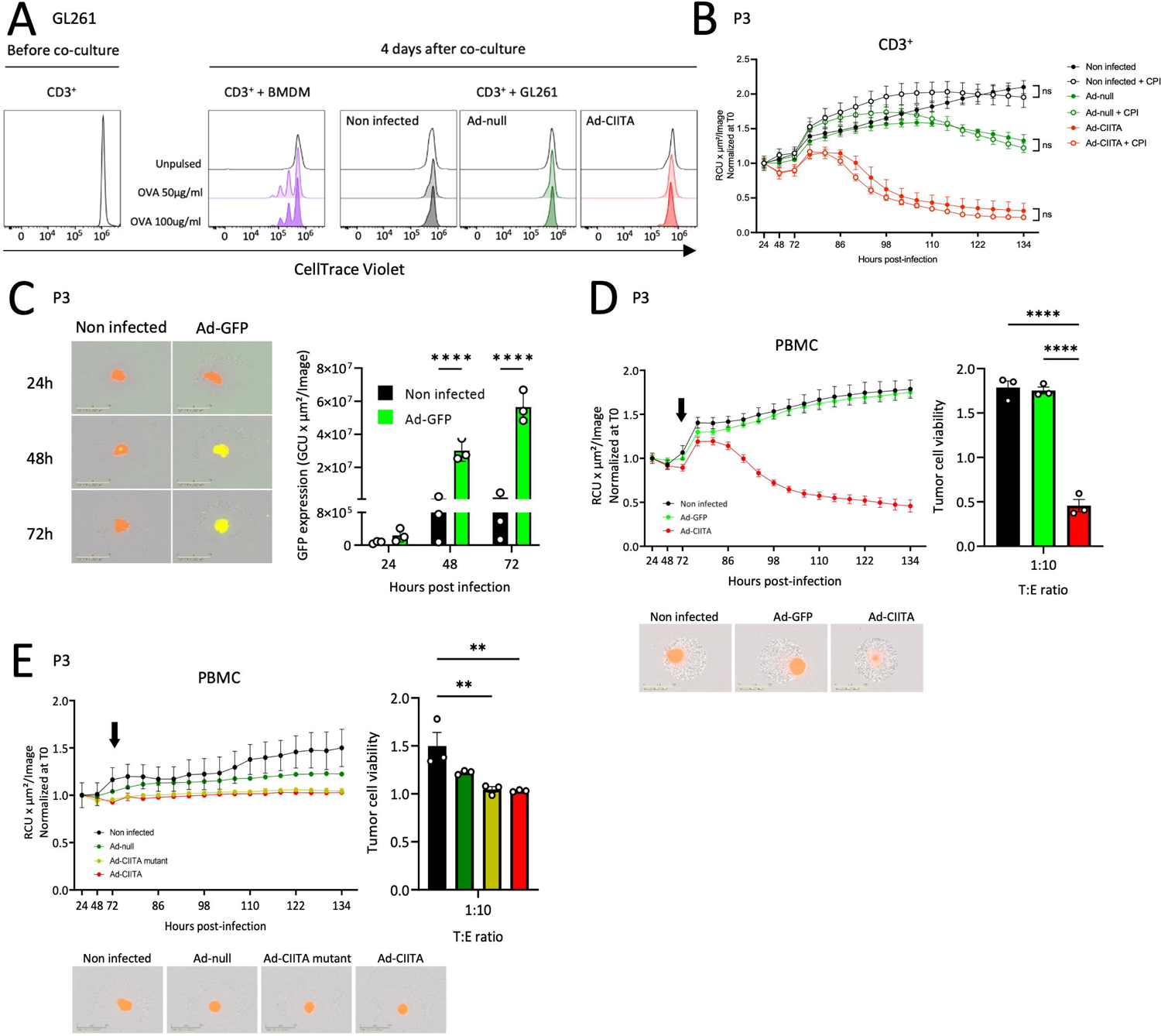
CIITA expression and cell-to-cell contact, but not antigen presentation, are required for immune-mediated tumor cell killing. **A.** *In vitro* proliferation assay of CD3^+^ T-cells co-cultured with OVA323-339-pulsed GL261 tumor cells or BMDM (positive control). **B.** Tumor cell viability in P3dsRed organoids (MOI 75) when co-cultured with CD3^+^ T-cells (T:E ratio of 1:10), comparing the effects of unpulsed versus CPI pool-pulsed organoids. **C-E.** P3dsRed organoids (MOI 75) co-cultured with PBMCs (T:E - 1:10). **C.** GFP expression in Ad-GFP-infected vs non-infected organoids. Mean ± SD. ****p<0.0001 (two-way ANOVA). **D.** Tumor cell viability in Ad-GFP- or Ad-CIITA-infected GB organoids compared to non-infected control. Mean ± SEM. ****p<0.0001 (one-way ANOVA). **E.** Tumor cell viability in two-layer transwell co-culture model. Mean ± SEM. **p<0.01 (one-way ANOVA).

To address the transgene specificity of the observed phenotype, we infected P3 organoids with an Ad-GFP vector (**Fig.5C**). Unlike Ad-CIITA, Ad-GFP did not lead to tumor cell death upon co-culture with PBMCs (**Fig.5D**). The same results were obtained in the T16 co-culture model (**Supplementary Fig.5A-B**), confirming the selectivity of the immune-mediated phenotype toward CIITA.

Next, we determined whether tumor killing required cell-to-cell contact by performing co-cultures in a two-layer transwell system. While we observed the expected reduction in organoid growth due to AdV infection, organoid breakdown in the presence of Ad-CIITA and Ad-CIITA mutant was completely abolished (**Fig.5E**). This was replicated in T16 organoids (**Supplementary Fig.5C**), indicating that immune cells require direct cell-to-cell interaction with target GBs to effectively induce tumor cell death.

### Immune-mediated tumor cell killing of GB organoids is not induced by degranulating T-cells or by activation of death receptor signaling pathways

We next aimed to assess different modes of contact-dependent T-cell-mediated cell killing. Cytotoxic T-cells conventionally release lytic granules, such as perforin and granzymes, into the immunological synapse. T-cell degranulation did not appear to be involved in CIITA-mediated killing, as evidenced by the absence of the degranulating marker CD107a and intracellular IFN-γ expression (**Fig.6A**). Cell death can also be induced via activation of death receptor signaling in target cells, including tumor necrosis factor alpha (TNF-α)/TNF-receptor (TNF-R) and Fas/Fas ligand (FasL)^18^. Blocking these death pathways via anti-human TNF-R and FasL monoclonal antibodies did not impact the ability of CIITA to induce GB organoid killing (**Fig.6B**), in contrast to Actinomycin D- and dimethyl sulfoxide (DMSO)-induced cell killing, respectively (**Supplementary Fig.6**).

**Figure 6.**
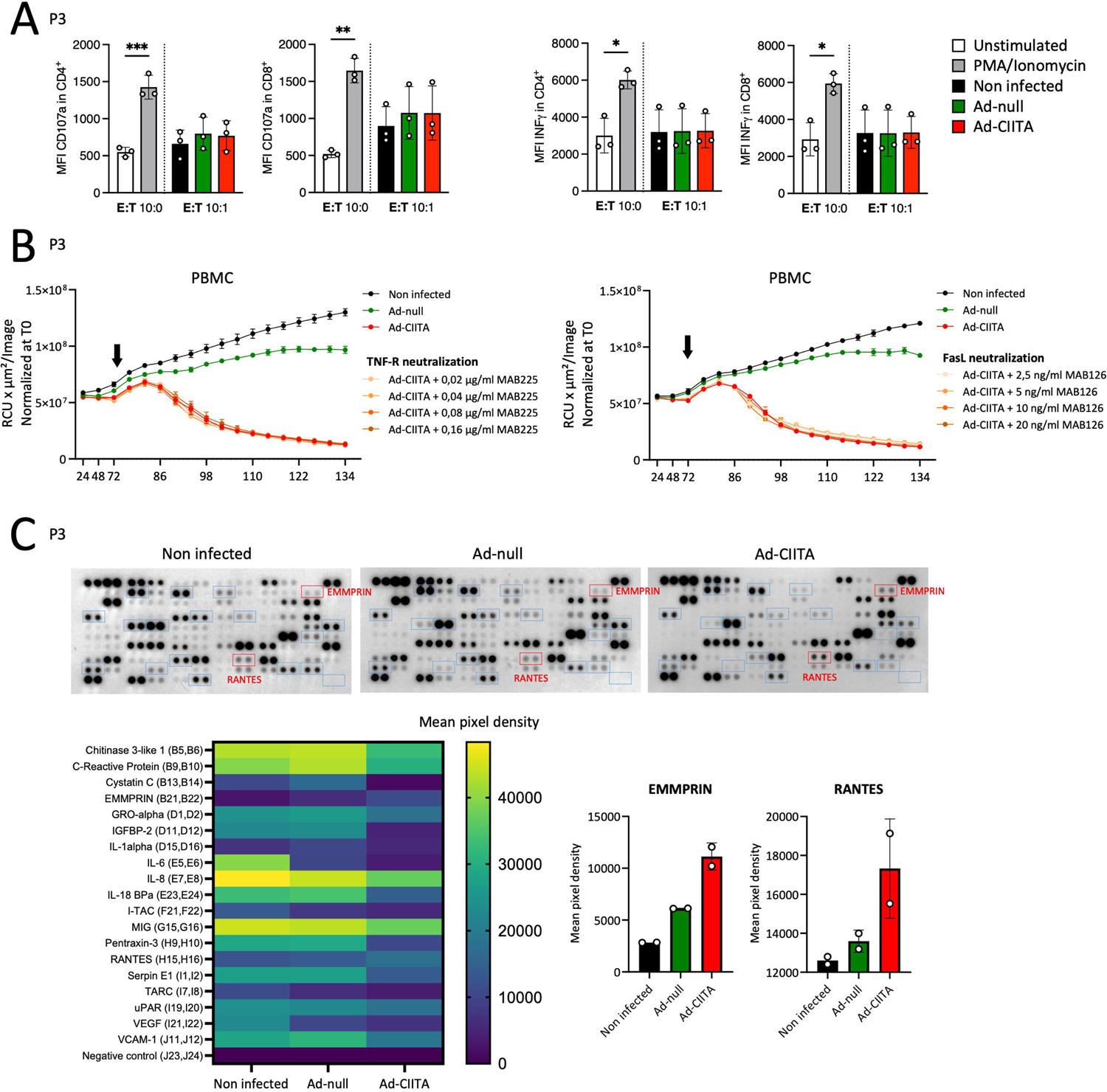
Immune-mediated tumor cell killing of GB organoids does not involve T-cell degranulation or canonical death receptor signaling pathways. **A.** FACS analysis of surface CD107a (left panel) and intracellular IFN-γ (right panel) in CD4^+^ and CD8^+^ T-cells at 24 hours post co-culture. Mean ± SD. *p<0.05, **p<0.01 and ***p<0.001 (one-way ANOVA). **B.** Neutralization assay using anti-human TNF-R (MAB225) (left panel) or anti-human FasL (MAB126) (right panel). Mean ± SEM. **C.** Heatmap of the most upregulated and downregulated proteins detected via protein cytokine array (bottom left). Graphs showing levels of secreted EMMPRIN and RANTES (bottom right). Mean ± SD.

In an attempt to identify non-classical T-cell activation mechanisms, we investigated the secretion of cytokines upon tumor cell killing mediated by wild-type CIITA at 72 hours post co-culture (**Fig.6C**). We detected increased secretion of the extracellular matrix metalloproteinase inducer (EMMPRIN) and RANTES (Regulated upon Activation, Normal T-cell Expressed, and Secreted) in Ad-CIITA co-culture medium and a decrease in the levels of IL-1alpha, IL-18, VEGF, and IL-6. To what extent such changes in cytokine levels are functionally linked to tumor cell killing remains to be determined.

## DISCUSSION

CIITA is the master regulator of MHC-II expression, and this biological function has been exploited to trigger adaptive immunity against tumors^19^. Given the need to develop new strategies against GB tumors, we equipped a non-lytic AdV with the wild-type *CIITA* transgene (Ad-CIITA). At basal level, GB tumor cells generally lacked MHC-II expression, while other components of the MHC-II transcriptional machinery were present. Indeed Ad-CIITA infection of primary GB cells resulted in robust MHC-II expression at the surface of infected cells.

We found that while Ad-CIITA infection has limited impact on tumor cell growth, effective killing of infected GB organoids is activated and maintained only in the presence of immune cells. Our study reveals a new ability of CIITA to activate T-cells which is independent of MHC-II molecules and antigen presentation. Firstly, we find that a novel CIITA mutant, unable to induce MHC-II still triggers T-cell mediated tumor cell death. Secondly, depletion of either CD4^+^ or CD8^+^ cells did not impair immune-mediated cell death. Thirdly, cell killing was independent of antigen presentation. A direct influence of the virus could be ruled out by the use of control empty and GFP-containing vectors. Cell-to-cell contact between target GB and immune cells was necessary, yet, we did not detect evidence of canonical T-cell degranulation or known death signaling pathways. The mechanism by which CIITA induces an MHC-II independent immune-mediated tumor cell death remains to be determined.

The newly generated *CIITA* mutant harbors a point mutation (G1088A) within the carboxy-terminal region (4^th^ LRR domain) of an otherwise full-length protein. The replacement of the non-polar, small amino acid glycine with arginine, may impact protein folding, stability and binding properties^20^. While wild-type CIITA protein is distributed between the nucleus and cytoplasm^7^, the G1088A mutant protein was effectively excluded from the nucleus, in line with a previous report^21^. Our novel point-mutated CIITA variant may thus effectively serve to unravel MHC-II-dependent and - independent activities of CIITA. Cytoplasmic effects of CIITA have been previously reported such as the L1035P CIITA mutant enhancing human immunodeficiency virion maturation through Gag processing by the viral protease^22^.

The cytotoxicity induced by both Ad-CIITA and Ad-CIITA mutant in patient-derived GB cells was primarily mediated by CD3^+^ T-cells, in line with *in vivo* results in mice^12, 13,11^. Nevertheless, an involvement of other immune components, such as natural killer cells or macrophages, cannot be currently excluded. Further investigation should also be directed in validating the anticancer role of wild-type and mutant CIITA in humanized GB models.

T-cell activation and proliferation commonly require at least two key events: (i) T-cell receptor (TCR) binding to a specific MHC-restricted antigen, and (ii) the interaction of costimulatory molecules on the surface of T-cells and APCs. We have shown that the killing phenotype relies on direct cell-to-cell interaction between the target tumor organoids and the immune cells and was observed despite the absence of CD80 and CD86 costimulatory molecules, which is in line with previous reports^8, 12, 13^.

A protein cytokine array revealed increased secretion of EMMPRIN and RANTES along with a decrease in various pro-inflammatory cytokines in Ad-CIITA co-culture medium. EMPPRIN is a glycoprotein expressed and secreted by various cell types, including lymphocytes and tumor cells, involved in metalloproteinase induction, energy metabolism, T-cell activation, co-stimulation, and proliferation^23^. RANTES (CCL5) is a potent proinflammatory chemokine that acts as a chemoattractant or immune cell activator^24^. RANTES-induced T-cell activation has various effects, including lymphocyte proliferation or apoptotic cell death, as well as the release of pro-inflammatory cytokines^24^. Further studies are warranted to fully understand the implications of soluble EMMPRIN and RANTES in the observed T-cell-mediated killing.

In conclusion, our findings highlight the potential of viral delivery of the *CIITA* transgene in human GB as a promising immunotherapeutic approach for enhancing adaptive responses against GB. The observed T-cell-mediated cytotoxicity, which appears to be largely independent of MHC-II antigen presentation, opens up new avenues for CIITA analysis beyond its canonical transcriptional activity.

## MATHERIALS AND METHODS

### Patient-derived preclinical models and cell culture

PDOX models, PDOX-derived *ex vivo* organoids, and PDOX-derived GSC lines were generated as described in^25^ ^26^. DsRed-expressing organoids were created as described in^27^. BG5, BG7, GG16, GG6, U251, and U87 were kindly provided by Prof. Rolf Bjerkvig, University of Bergen; NCH421, NCH465, and NCH644 by Prof. Christel Herold-Mende, University of Heidelberg; and GL261 by Prof. Roberto Accolla, University of Insubria.

### HLA typing

gDNA extracted from GB organoids with the DNeasy Blood & Tissue Kit (Qiagen, Cat# 69504), was sequenced at 11 HLA loci (MHC-I *HLA-A*, *-B*, *-C*; MHC-II *HLA-DRB1/3/4/5*, *-DQA1*, *-DQB1*, *-DPA1*, *-DPB1*) using the Illumina TruSight HLA v2 Sequencing panel (**Table S3**).

### Gene expression analysis

The expression of CIITA and MHC-II associated genes in GB patient tumors and control normal brain was investigated with GEPIA2^28^. To examine gene expression in single cells isolated from GB patient tumors, we exploited the GBap dataset^29^ and the DotPlot function of Seurat package. Gene expression in preclinical models was assessed with the Gene-Chip® Human Gene 1.0 ST Arrays (Affimetrix, Cat# 901086) as described previously^26^ or RNA-seq on bulk tissue (50 million reads/sample, 2×150bp), using the Illumina Stranded mRNA prep kit on an Illumina NovaSeq 6000 sequencer.

### Peripheral blood lymphocytes

Allogeneic PBMCs were isolated from healthy human donor blood (Luxembourg Red Cross) via density gradient centrifugation and cryopreserved. Allogeneic HLA-matched PBMCs were purchased from Immunospot based on their HLA matching (**Table S3**). T-cell populations were isolated from PBMCs using CD3^+^ (BioLegend, Cat# 480022), CD4^+^ (Miltenyi, Cat# 130-096-533) or CD8^+^ (Miltenyi, Cat# 130-096-495) negative isolation kits. PBMC composition and purity were evaluated using multi-color flow cytometry (**Table S2**).

Immune cells were cultured in “T-cell medium”: RPMI 1640 GlutaMAX™ (Thermo Fisher, Cat#61870036), 9% FBS, 1% human serum (Sigma-Aldrich, Cat#H3667), 0.1 mM Sodium Pyruvate (Thermo Fisher, Cat#11360070), 55 μM β-Mercaptoethanol (Thermo Fisher, Cat#21985023), 1X NEAAs, 5 mM HEPES (Thermo Fisher, Cat#15630080), and 100 U/mL penicillin‒streptomycin.

### AdV vectors

Replication-deficient (ΔE1-E3) serotype 5 AdV vectors were created according to the Ad-Easy-1 protocol^30^. The AdVs used in the study include Ad-null (no transgene), Ad-GFP, Ad-CIITA (wild-type *CIITA* encoding for MHC-II TransActivator isoform III [NP_000237.2, 1130 aa]), and Ad-CIITA mutant (mutated version of *CIITA*). For all vectors, transgene expression was regulated via the CMV promoter. AdV particles were purified with the Adeno-X™ Mega Purification Kit (Takara, Cat# 631032) and titrated using the Adeno-X Rapid Titer kit (Takara, Cat# 632250), following manufacturer’s instructions.

AdV infection was conducted during cell seeding. After infection, cells were gently shaken at 37°C for up to one hour, and then incubated at 37°C for a maximum of six days post-infection, depending on the read-out.

### AdV-infected GB organoid - immune cell co-culture

Cryopreserved GB single cells were seeded (2E10^4^ cells/well) in 96-well, Nunclon Sphera-Treated, U-Shaped-Bottom, ultra-low attachment microplates (Thermo Fisher, Cat# 174929), infected in a final volume of 100 μL organoid medium, and incubated at 37°C on an orbital shaker at 40-50 rpm. Three days post-seeding/infection, cryopreserved immune cells were added at different T:E ratios (1:1, 1:5, 1:10). Before co-culture, GB organoids were pulsed with “CPI pool” (ImmunoSpot, Cat# CTL-CPI-001) as a positive control for T-cell activation. Co-cultures were performed in the presence of 25 U/mL IL-2 (Miltenyi, Cat# 130-097-745) and 5 μg/mL anti-CD28 antibody (BioLegend, Cat# 302902). Tumor cell viability was monitored in real-time using IncucyteS3, or via flow cytometry at 72 hours post-co-culture via staining of CD45^-^/CD3^-^/ CD90^+^ tumor cells with Live/Dead fixable Near-IR (**Table S4**). To study contact-dependency, co-cultures were carried out using transwell-96 with 0.4 µm pore polyester membrane inserts (StemCell Technologies, Cat# 100-0419).

### Incucyte S3 live cell imaging system

Incucyte S3 (RRID:SCR_023147) was used to monitor tumor cell viability over time (4X magnification). Organoids were imaged daily for the first 3 days post-seeding/infection and every 4 hours after co-culture with immune cells. Data were analyzed using Incucyte 2022A software.

To quantify tumor cell viability in dsRed-expressing organoids (P3dsRed and T16dsRed), we monitored total integrated red fluorescence intensity (RCU x µm²/Image) over time. For T188 tumor cells transiently labeled with 0.3 μM of Incucyte® Cytolight Rapid Red Dye (Sartorius, Cat# 4706), we measured the red total area (μm^2^/Image) over time.

### Degranulation assay

At 24 hours post co-culture (T:E ratio of 1:10), CD3^+^ T-cells were labelled with anti-CD107a antibody (**Table S4**). Freshly thawed CD3^+^ T-cells stimulated with 50 ng/mL phorbol 12-myristate 13-acetate (PMA) (Enzo Life Sciences, Cat# BML-PE160-0001) and 1 µg/mL ionomycin (Enzo Life Sciences, Cat# ALX-450-007-M001) were used as positive control. Cells were then incubated for 4 hours at 37°C with GolgiStop™ (BD, Cat#554724, 1:1000 dilution), and GolgiPlug (BD, Cat#555029, 1:500 dilution). CD3^+^ T-cells from five different wells (∼1 million cells) were pooled together, stained with cell surface markers for 30 minutes at 4°C, fixed/permeabilized with Cytofix/Cytoperm (BD, Cat#554714) for 20 min at 4°C, and finally incubated with anti-IFN-γ intracellular antibody for 30 minutes at 4°C (**Table S4**).

### *In vitro* T-cell proliferation assay

GL261 cells were infected and pulsed with two concentrations (50 µg/ml and 100 µg/ml) of OVA323-339 (Eurogentec, Cat#AS27024) for 4 hours at 37°C. After extensive washes, they were co-cultured with Cell Trace Violet (Thermo Fisher, Cat#C34557)-stained CD3+ T-cells from OT-II mice. As a positive control, bone marrow-derived macrophages were also pulsed with OVA323-339 and co-cultured with OT-II CD3+ T-cells. CD3+ T-cells were acquired using the NovoCyte Quanteon Flow Cytometer on day 0 (before co-culture) and on day 4 (four days post-co-culture) to trace CD4^+^ T-cell proliferation.

### Neutralization assay

Organoids were treated with increasing concentrations of anti-human TNF-RI/TNFRSF1A (R&D, Cat# MAB225) (0.16 μg/mL, 0.08 μg/mL, 0.04 μg/mL, and 0.02 μg/mL) and anti-human Fas Ligand/TNFSF6 (R&D, Cat# MAB126) (20 ng/mL, 10 ng/mL, 5 ng/mL, and 2.5 ng/mL) monoclonal antibodies (**Table S4**). Organoids treated with actinomycin D (1 μg/mL) or 2.5% DMSO were used as positive controls for TNF-R and FasL neutralization, respectively. Immune cell-mediated tumor killing was assessed via Incucyte S3.

### Measurement of cytokine secretion

At 72 hours post PBMC co-culture (T:E ratio of 1:10), conditioned media were collected, centrifuged, and screened using the Proteome Profiler Human XL Cytokine Array Kit (R&D, Cat# ARY022B). Array membranes were then exposed to X-Ray films (1, 5, and 10 minutes). The average signal (pixel density) per analyte was quantified using Quick Spots software.

### Quantitative RT‒PCR

Total RNA was extracted using the RNeasy Mini Kit (Qiagen, Cat# 74104), reverse-transcribed to cDNA using Fast SYBR Green (Thermo Fisher, Cat# 4385612), and QuantStudio 5 Real-Time PCR System (RRID:SCR_020240). Relative gene expression levels were calculated through the ΔCT method by using *elongation factor 1 alpha (EF1α)* as a housekeeping gene. The primers used were as follows: CIITA (Fw: TTATGCCAATATCGCGGAACTG, Rv: CATCTGGTCCTATGTGCTTGAAA), HLA-DRα (Fw: AGTCCCTGTGCTAGGATTTTTCA,--Rv: ACATAAACTCGCCTGATTGGTC),-EF1α-(Fw: TTGTCGTCATTGGACACGTAG,-Rv:-TGCCACCGCATTTATAGATCAG), and-GAPDH-(Fw: CTCCCACTCTTCCACCTTCG, Rv: CCACCACCCTGTTGCTGTAG).

### Immunofluorescence

Cells were seeded/infected (5E10^4^ cells/well) in 8-well µ-slides (Ibidi, Cat# 80826) and, three days post-infection, were fixed with 4% PFA for 15 minutes at RT and blocked in 1X PBS with 0.5% Triton X-100 (Thermo Fisher, Cat# 85111) and 10% FBS for 1 hour at RT. Cells were then incubated with anti-CIITA primary antibody (Abcam, Cat# ab92511) diluted in 1X PBS with 0.5% Triton X-100 overnight at 4°C (**Table S4**). After three washes in 1X PBS, the cells were incubated with Alexa Fluor 488 secondary antibody (Thermo Fisher, Cat# A11001) and Hoechst dye (Thermo Fisher, Cat# H3569) diluted in 1X PBS with 0.01% BSA (Sigma-Aldrich, Cat# A9418) for 1 hour at RT (**Table S4**). Cells were then imaged with a Zeiss LSM 880 Confocal Laser Scanning Microscope (RRID:SCR_020925) and Zen Blue software (RRID:SCR_013672).

### Flow cytometry

Single cells were stained with fluorochrome-conjugated antibodies (**Table S4**) diluted in ice-cold 1X PBS with 0,2% BSA (“FACS buffer”) or Brilliant Stain Buffer (“BV buffer”) (BD, Cat# 563794) for 30 minutes at 4°C in the dark. Stained cells (1 million cells in 100 μL, we scaled down accordingly for a smaller number of cells) were acquired with NovoCyte Quanteon Flow Cytometer. Data were analyzed using FlowJo (RRID:SCR_008520) version 10.

### Statistical analysis

Experimental results were analyzed using GraphPad Prism 9 (RRID:SCR_002798) and reported as mean ± standard deviation (SD) or, for co-culture experiments, as mean ± standard error of mean (SEM). For flow cytometry results, figures depict expression curves, along with the mean percentage of positive cells and their standard deviation (based on three independent experiments). All means were calculated based on a minimum of three independent experiments. For co-culture experiments, all means were calculated based on a minimum of three immune cell donors, representing three different biological replicates. One-way ANOVA was employed to compare multiple groups, while two-way ANOVA was used to compare two or more factors simultaneously. A P value < 0.05 was used to determine statistical significance.

## ACKNOWLEDGEMENTS

We are grateful for the use of the LIH PRECISION-PDX Brain Tumor Bank (www.precision-pdx.lu), supported by the patients, the Neurosurgery Department of the Centre Hospitalier de Luxembourg, the NORLUX Neuro-Oncology Laboratory and the Clinical and Epidemiological Investigation Center of LIH. We thank Prof. Roberto Accolla and Prof. Greta Forlani for providing cell lines and critically reviewing the manuscript, Pilar M. Moreno-Sanchez for technical assistance in the management of PDOXs, and the Luxembourg Croix Rouge and its donors for generously providing healthy donor blood samples. We acknowledge the support from LIH’s core facilities (Animal facility, In vivo imaging facility, National Cytometry Platform, LUXGEN sequencing platform).

## AUTHOR CONTRIBUTION

Conceptualization (I. S., A.P. A.G., S.P.N., A.M.), methodology (I.S., E.K., A.P., M.R., A.C., A.G.), investigation (I.S., E.K., A.P., M.R., A.L., A.O., V.B., G.M.D.), formal analysis (L.E., B.N., A.G.), supervision (A.P., A.G., A.M., S.P.N.) writing - original draft (I.S., A.G., A.M., S.P.N.), revision (I.S., E.K., A.G., S.P.N.) – review & editing (all authors). The authors approved the final version of the manuscript as submitted and agreed to be accountable for all the aspects of the work.

## CONFLICT OF INTEREST

The authors have nothing to disclose.

## FUNDING INFORMATION

We acknowledge the financial support by the Luxembourg Institute of Health, FNRS-Télévie (GBodImm no. 7.8513.18, GBodImm2 no. 7651720F, ImmoGB 7.8505.20, ImmoGB2 7.6603.22) and the Luxembourg National Research Fund (C20/BM/14646004/GLASSLUX, C21/BM/15739125/DIOMEDES).

## DATA AVAILABILITY STATEMENT

The authors confirm that the data supporting the findings of this study are available within the article and its supplementary materials.

## SUPPLEMENTARY FIGURES

**Supplementary Figure 1.**
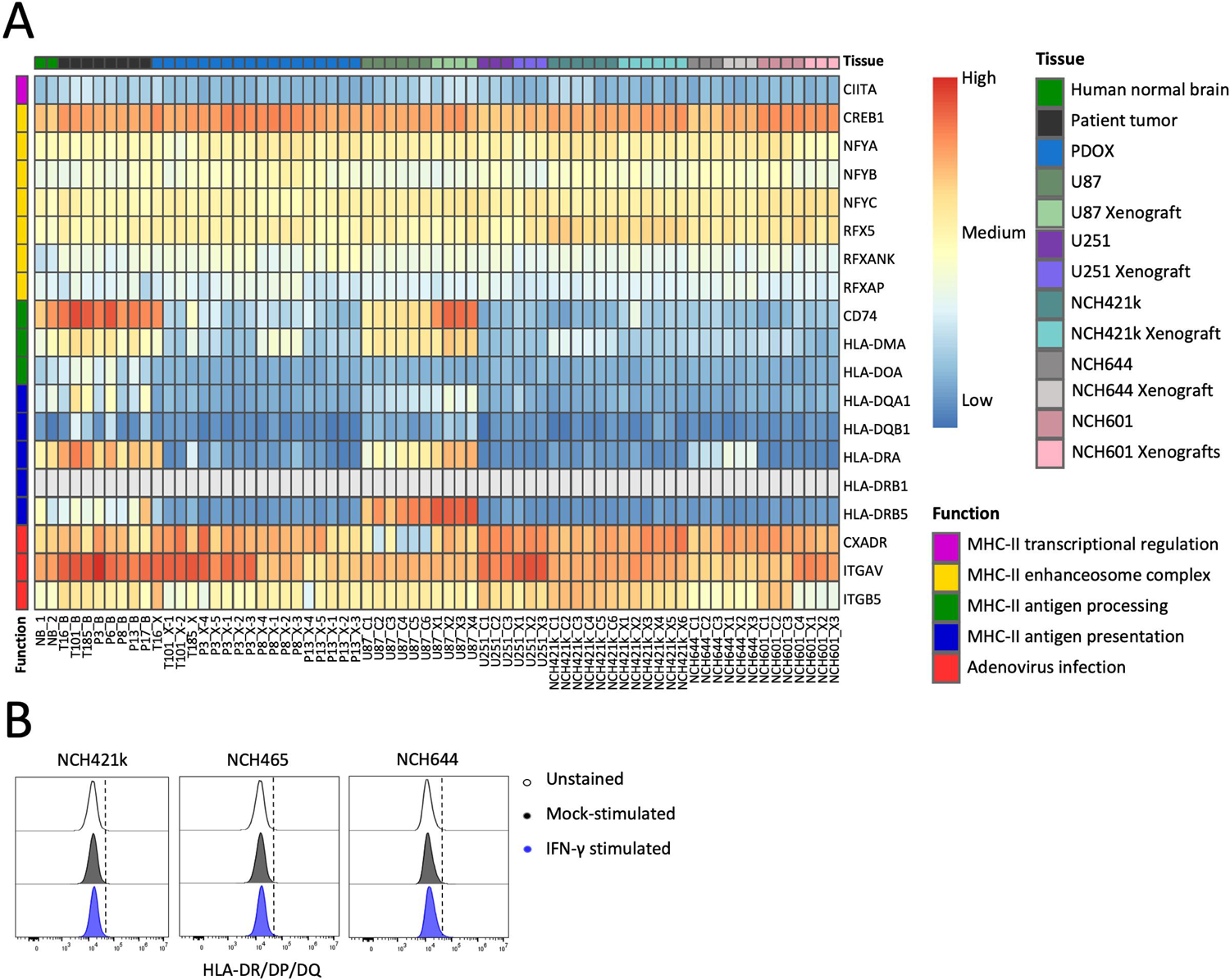
MHC-II related gene expression analysis in GB patient tumors and human preclinical models. **A.** Heatmap representing gene expression levels in normal human brain, GB patient tumors, PDOXs, GSC lines (NCH421k, NCH644), and classical glioma lines (U87, U251) grown *in vitro* or *in vivo* as xenograft (’X’). Lack of expression is represented by grey color. Human specific arrays were applied for transcriptome analysis. **B.** FACS analysis of surface MHC-II (anti-HLA-DR/DP/DQ-FITC) in control conditions (cells unstained or mock-stimulated) and upon IFN-γ stimulation.

**Supplementary Figure 2.**
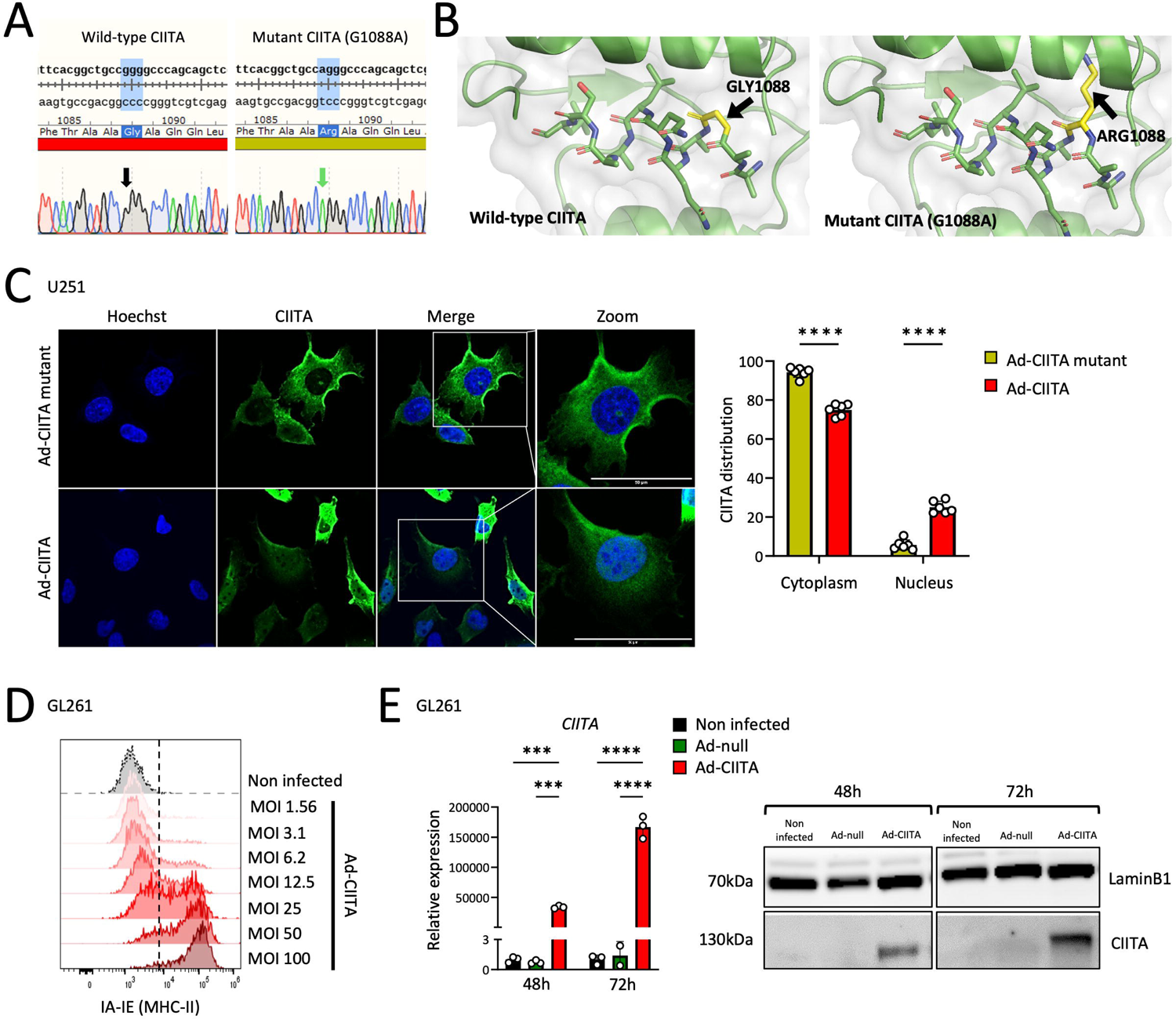
Characterization of wild-type and mutant CIITA adenoviral vectors in adherent human (U251) and murine (GL261) cell lines. **A.** Sanger sequencing results of the wild-type vs mutant *CIITA* DNA. **B.** PyMOL representation of the amino acid string surrounding position 1088 in wild-type (left) and mutated (right) CIITA. **C.** Representative confocal images of CIITA (Alexa Fluor 488) in U251 at 72 hours post-infection (magnification 63X). CIITA cellular distribution was quantified by measuring the ratio of cytoplasmic-to-nuclear fluorescence intensity. Mean ± SD. ****p<0.0001 (two-way ANOVA). **D-F.** Ad-CIITA characterization in GL261. **D.** FACS analysis of MHC-II (anti-IA-IE-PerCP) at increasing Ad-CIITA MOIs. **E.** mRNA (left) and protein (right) levels of CIITA at 48 and 72 hours post-infection, as analyzed by qRT‒PCR and Western blot, respectively. qRT-PCR and WB results were normalized to GAPDH. Mean ± SD. ***p<0.001, ****p<0.0001 (two-way ANOVA).

**Supplementary Figure 3.**
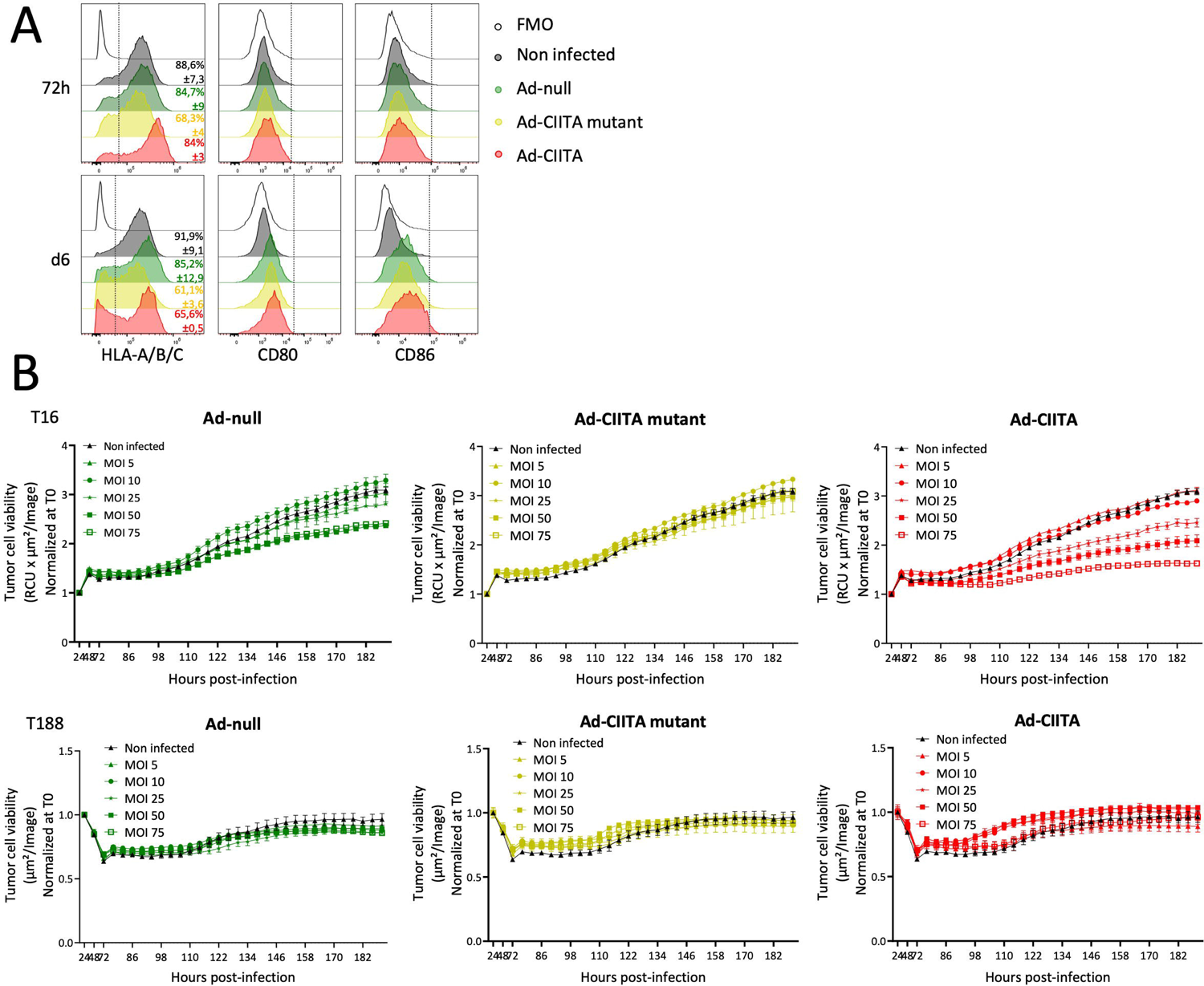
Infection of human primary glioblastoma organoids with adenoviral vectors and impact on MHC-I expression. **A.** FACS analysis of surface MHC-I (anti-HLA-A/B/C-BV510), CD80 (anti-CD80-PE) and CD86 (anti-CD86-BV421) in P3 organoids. **B.** Tumor cell viability of T16dsRed (top) and red dye-labeled T188 (bottom) organoids infected at increasing virus MOIs. Mean ± SEM.

**Supplementary Figure 4.**
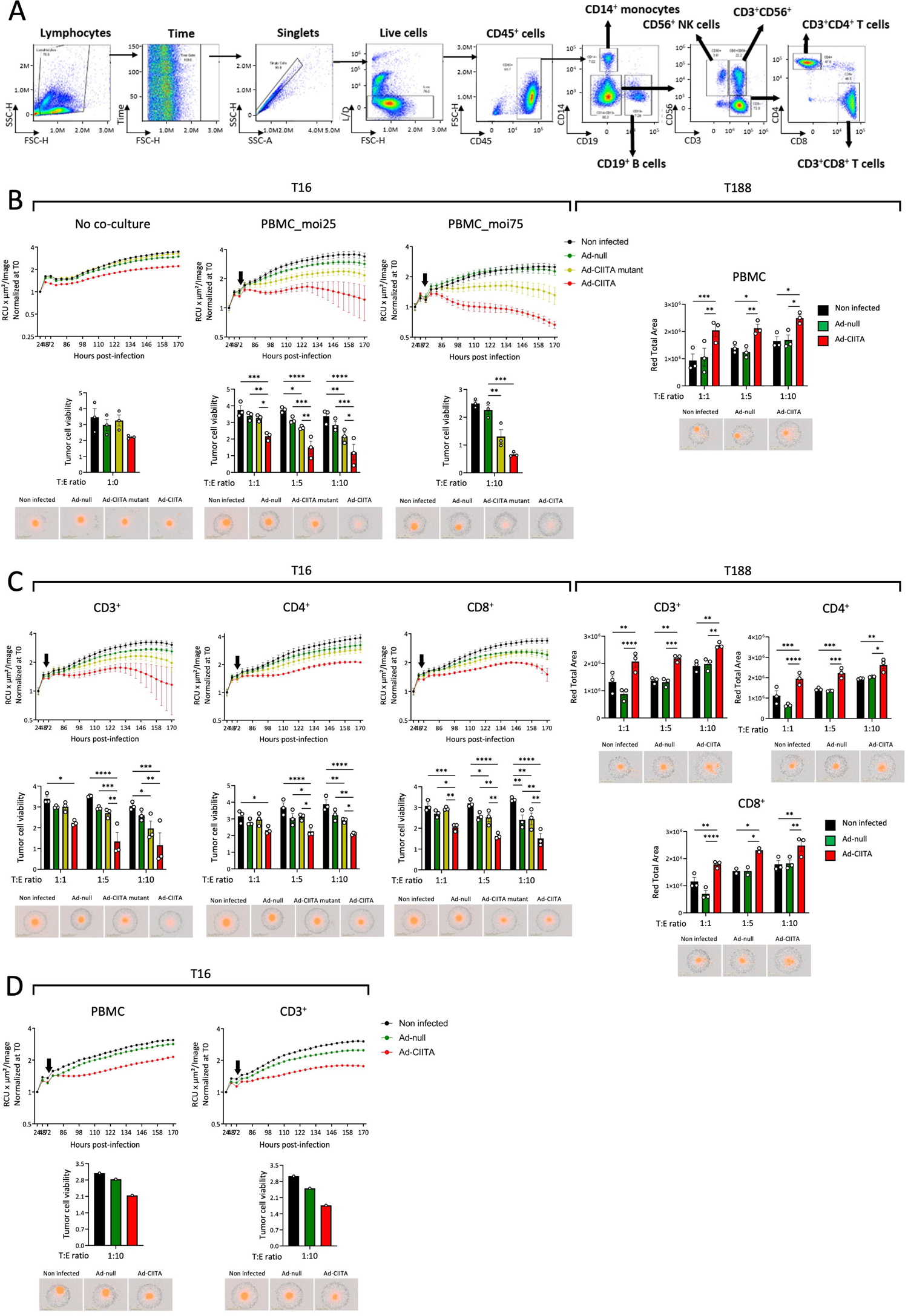
Evaluation of immune cell-mediated tumor cell killing in human primary GB organoids. **A.** FACS gating strategy to phenotype PBMCs and MACS-purified T-cell fractions. **B-C.** Tumor cell viability in T16dsRed (left panel, MOI 25) or red dye-labeled T188 (right panel, MOI 75) organoids: **(B)** alone (no co-culture) or co-cultured with PBMCs; **(C)** co-cultured with CD3^+^, CD4^+^, or CD8^+^ T-cells. **D.** Tumor cell viability in T16dsRed organoids (MOI 25) co-cultured with HLA-matched PBMCs or CD3^+^ T-cells. Mean ± SEM. *p<0.05, **p<0.01, ***p<0.001, and ****p<0.0001 (two-way ANOVA). Deviations from virus MOI are explicitly indicated on the graph.

**Supplementary Figure 5.**
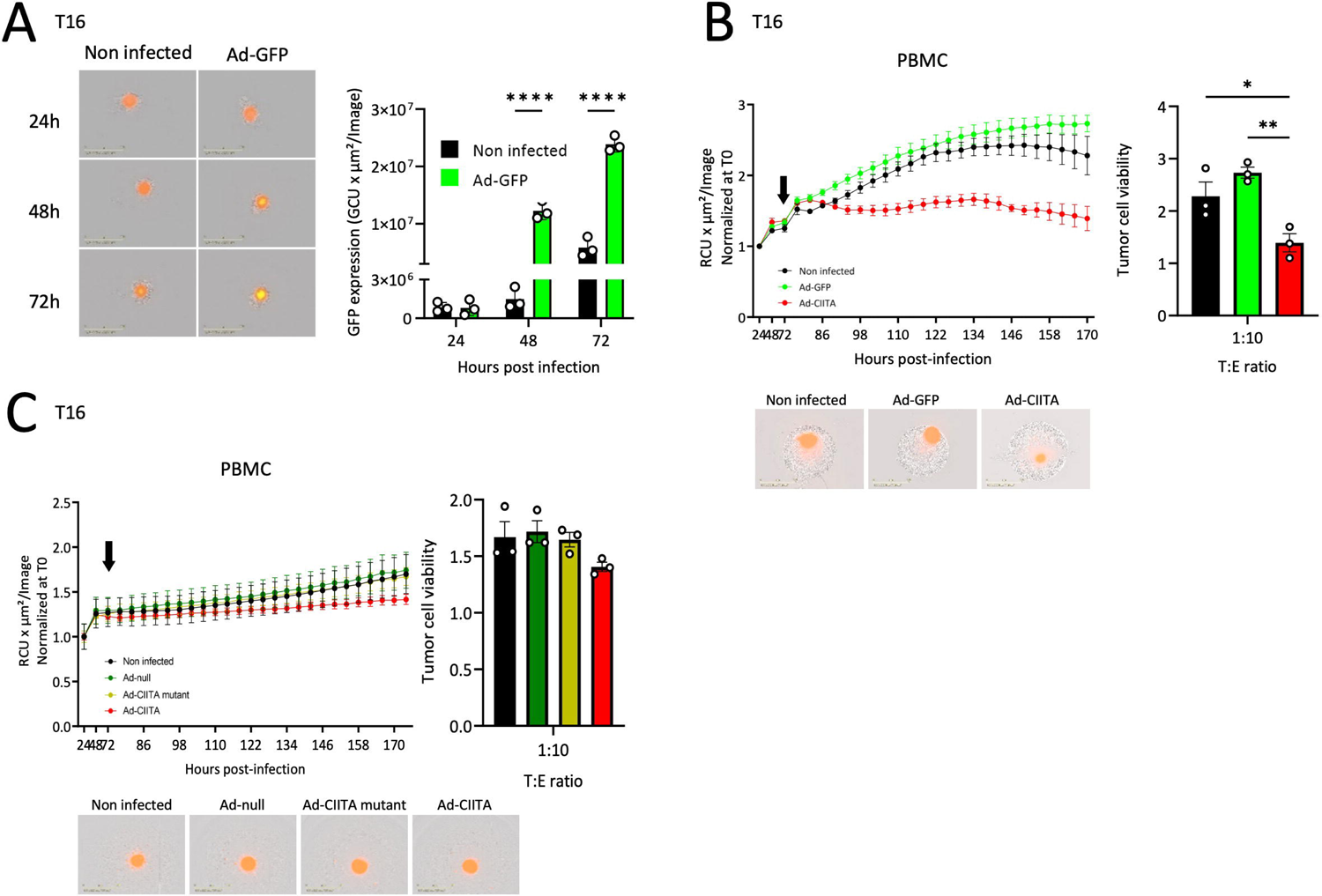
Requirement for CIITA expression and immune-tumor cell contact, but not antigen presentation. **A-C.** T16dsRed organoids (MOI 25) co-cultured with PBMCs (T:E - 1:10). **A.** GFP expression in Ad-GFP-infected vs non-infected organoids. Mean ± SD. ****p<0.0001 (two-way ANOVA). **B.** Tumor cell viability in Ad-GFP- or Ad-CIITA-infected GB organoids compared to non-infected control. Mean ± SEM. *p<0.05, **p<0.01 (one-way ANOVA). **C.** Tumor cell viability in two-layer transwell co-culture model. Mean ± SEM.

**Supplementary Figure 6.**
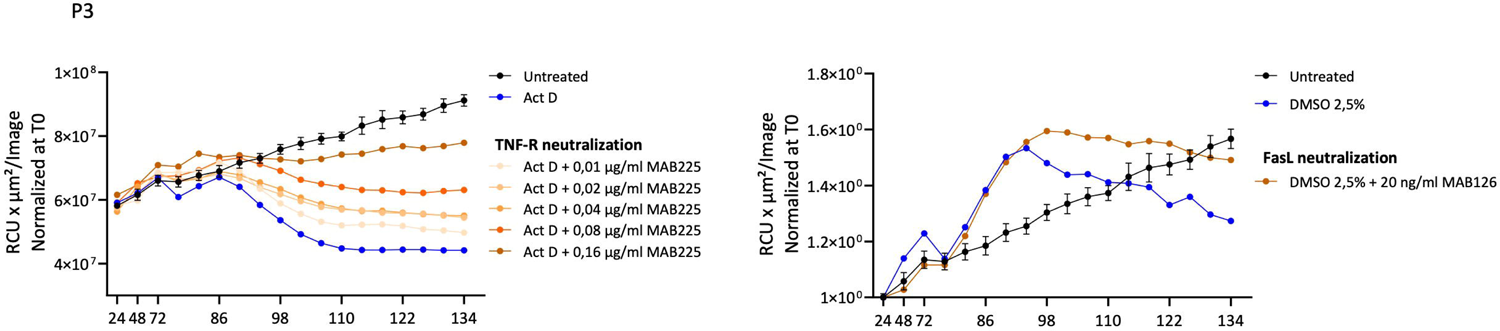
Positive controls of neutralization assay. Actinomycin D- and DMSO-induced tumor cell death blocked by anti-human TNF-R (MAB225) (left panel) and anti-human FasL (MAB126) (right panel), respectively. Mean ± SEM.

## Notes

### Competing Interest Statement

The authors have declared no competing interest.

### Summary of Updates

This version of the manuscript has been revised to enhance clarity and quality. The word count has been significantly reduced by eliminating redundant information and streamlining the discussion. Data presentation has been improved by updating figures and tables. New figures have been added, and some supplementary data has been integrated into the main text for a more comprehensive understanding. References have been updated to include recent developments in the field.

